# Kaminari: a resource-frugal index for approximate colored *k*-mer queries

**DOI:** 10.1101/2025.05.16.654317

**Authors:** Victor Levallois, Yoshihiro Shibuya, Bertrand Le Gal, Rob Patro, Pierre Peterlongo, Giulio Ermanno Pibiri

## Abstract

**Motivation:** The problem of identifying the set of textual documents from a given database containing a query string has been studied in various fields of computing, e.g., in Information Retrieval, Databases, and Computational Biology. We consider the approximate version of this problem, that is, the result set is allowed to contain some false positive matches (but no false negatives), and focus on the specific case where the indexed documents are DNA strings. In this setting, state-of-the-art solutions rely on Bloom filters as a way to index all *k*-mers (substrings of length *k*) in the documents. To answer a query, the *k*-mers of the query string are tested for membership against the index and documents that contain at least a user-prescribed fraction of them (e.g., 75–80%) are returned.

**Methods and results:** Here, we explore an alternative index design based on *k*-mer minimizers and integer compression methods. We show that a careful implementation of this design outperforms previous solutions based on Bloom filters by a wide margin: the index has lower memory footprint and faster query times, while false positive matches have only a minor impact on the ranking of the documents reported. This trend is robust across genomic datasets of different complexity and query workloads.

**Software:** The software is implemented in C++17 and available under the MIT license at github.com/yhhshb/kaminari. Reproducibility information and additional results are provided at github.com/vicLeva/benchmarks_kaminari.

## 1. Introduction

Let ℛ = {*R*_1_, …, *R*_*N*_} be a collection of textual documents. Efficiently identifying which documents in ℛ contain a query string *Q* is a fundamental problem, especially in fields like Information Retrieval and Computational Biology. This work focuses on a specialized case where the documents are DNA strings, using the alphabet Σ = {A, C, G, T}. For large-scale processing, it has become customary to represent a DNA string by its collection of *k*-long substrings (named “*k*-mers”). To assess whether *Q* is present in a document *R*_*i*_, we compute the number of *k*-mers of *Q* that are also substrings of *R*_*i*_. If this number is at least 75–80% of the total *k*-mers in *Q*, we consider *Q* to appear in *R*_*i*_ [Ukkonen, 1992]. Therefore, *R*_*i*_ can be viewed as a (multi-)set of |*R*_*i*_| − *k* + 1 *k*-mers. However, *k*-mers of *Q* may appear in different order in *R*_*i*_, which would mean *Q* does not actually appear as a substring. This situation is unlikely unless *k* is very small. In this work we study efficient solutions to the following problems.

### Problem 1

(Exact colored *k*-mer indexing.) Build a data structure, referred to as the *index* in the following, that allows to retrieve the set *C*_*k*_(*x*) = {*i*|*x* ∈ *R*_*i*_} as efficiently as possible for any *k*-mer *x* ∈ Σ^*k*^. If *x* does not occur in any *R*_*i*_, then *C*_*k*_(*x*) = ∅.

In other words, the set *C*_*k*_(*x*) contains all the identifiers, called “colors”, of the documents where the *k*-mer *x* appears. We refer to *C*_*k*_(*x*) as the *color set* of *x*.

While exact solutions to Problem 1 have been studied extensively [Karasikov et al., 2020, 2022; Alanko et al., 2023; Fan et al., 2023, 2024], *approximate* solutions — allowing for potential *false positive* matches but no *false negatives* — are gaining importance for their potential enhanced performance and scalability. In this work, we thus consider the approximate version of this problem, defined as follows.

### Problem 2

(Approximate colored *k*-mer indexing.) Build an index allowing retrieval of set 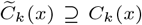 as efficiently as possible, for any *k*-mer *x* ∈ Σ^*k*^ and with small 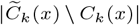.

The primary motivation for studying solutions to Problem 2 is to create a more efficient method for computing 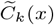, requiring less RAM and CPU time than *C*_*k*_(*x*). As a drawback, 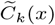 is a super set of *C*_*k*_(*x*), which may lead to false positives (elements in 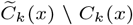 that do not actually contain the *k*-mer *x*.

State-of-the-art solutions to Problem 2 (reviewed in Section 3) are primarily based on *Bloom filters* [Bloom, 1970] — the most well-known approximate membership data structure. In these solutions, a Bloom filter is created for each document by inserting all its *k*-mers in the filter. The set of filters is then laid out as a matrix [Bingmann et al., 2019; Bradley et al., 2019; Lemane et al., 2022, 2024], or in a tree hierarchy [Mehringer et al., 2022; Marchet and Limasset, 2023; Sun et al., 2018; Harris and Medvedev, 2020]. As argued below, these approaches are space-inefficient and do not exploit some important properties of *k*-mers to obtain better performance for Problem 2.

### Our contribution

We contribute an alternative index design that does not use Bloom filters but is based on *k*-mer minimizers [Schleimer et al., 2003; Roberts et al., 2004] and integer compression techniques [Pibiri and Venturini, 2021] to address the main limitations of the state of the art.

In short, our solution merges the color sets of *k*-mers in a coherent way, e.g., those that share the same minimizer (smallest substring). This allows to save storage space (as the number of indexed sets reduces dramatically).Unlike Bloom-filter based solutions, where approximate color sets are created by merging the color sets of *k*-mers drawn at random (*k*-mers that hash to the same bit positions in the filter), we merge them based on their “similarity”.

While this merging strategy still allows false positives, they minimally affect the result sets computed by our index in the following sense. For a query *Q*, each reported document in the result set is associated to a *weight*, being the number of *k*-mers of *Q* that it contains. Retrieved documents are sorted by decreasing weight, allowing to rank results. We argue that most false positives do not have a weight able to *significantly alter the ranking of the result set* compared to an exact solution. To assess this result, we use the established similarity measure called *rank-biased overlap* [Webber et al., 2010].

These methods have been implemented in a software called “Kaminari” (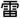, “thunder” in Japanese), freely available on GitHub. We conducted an extensive experimental analysis, comparing Kaminari with other efficient approximate and exact indexes. Results show that Kaminari produces smaller indexes and faster queries than other approaches. Kaminari is also competitive in terms of time and resources used to build the index. Lastly, we demonstrate that false positives have a small impact on the user, as the most relevant results of a query remain trustworthy.

## 2. Background

### 2.1. Sampling and hashing

#### Definition 1

(Minimizer sampling.) A minimizer sampling scheme is defined by a triple (*m, k*, 𝒪), with *m, k* ∈ℕ, *m < k*, and 𝒪 : Σ → ℝ^*m*^ is an order over all *m*-long strings. Given a *k*-mer *x*, the minimizer *µ* of *x* is the leftmost *m*-mer of *x* such that 𝒪(*µ*) ≤ 𝒪(*y*) for any other *m*-mer *y* of *x*.

In practice, 𝒪 is usually implemented as a non-cryptographic pseudo-random hash function, such as MurmurHash2 [Appleby, 2016], obtaining the so-called “random” minimizer scheme [Schleimer et al., 2003; Roberts et al., 2004]. (We however use the simple lexicographic order for ease of visualization when discussing the examples.) To simplify notation, “Minimizer(*x*)” designates the minimizer of the *k*-mer *x*, without specifying parameters *m* and 𝒪. Also, Minimizers(*Q*) is the (multi-)set of all the minimizers of the *k*-mers of the string *Q*.

A string is said *sampled* at the positions of the minimizers of its *k*-mers. For a string composed of *n* i.i.d. random characters and when *m* is sufficiently long, the expected number of distinct sampled positions is ≈ 2*/*(*k* − *m* + 2) · (*n* − *m* + 1) (see Theorem 3 by [Zheng et al., 2020] for details and the paper by [Groot Koerkamp and Pibiri, 2024] for a recent overview of sampling algorithms).

#### Definition 2

(Minimal perfect hash function (MPHF).) Let *f* : *U* → {1, …, *n*} for some universe set *U*. The function *f* is said to be a minimal perfect hash function for the set *S* ⊆ *U*, with |*S*| = *n*, if *f* (*x*) ≠ *f* (*y*) for all *x, y* ∈ *S, x* ≠ *y*.

In simpler words, a MPHF for the set *S* maps its *n* keys bijectively into the first *n* natural numbers. The theoretical space lower bound [Mehlhorn, 1982; Mairson, 1983] for representing a MPHF is *n* log_2_ *e* − *O*(log *n*) bits assuming |*U* | → ∞, which is approximately 1.443 bits per key for large *n*. Practical constructions with just 0.1% overhead on top of the lower bound have been proposed [Lehmann et al., 2025]. In this work, we use the PTHash data structure [Pibiri and Trani, 2023; Hermann et al., 2024], optimized for fast queries with a space usage of 2 − 3 bits/key.

### 2.2. Query mode

Given a multi-set *M*, we indicate with *w*(*i, M*) the multiplicity of element *i* in *M*. We colloquially refer to *w*(*i, M*) as the *weight* of *i* in *M*.

#### Definition 3

(Threshold-union query.) Let *Q* be a query string with |*Q*| ≥ *k* and *M* (*Q*) = ⊎_*x*∈*Q*_*C*_*k*_(*x*) be the multi-set union of the color sets for all the *k*-mers of *Q*. For a given 0 *< τ* ≤ 1, the threshold-union query computes the list *R*(*Q, τ*) that is the set {*i*|*w*(*i, M* (*Q*)) ≥ ⌊*τ* (|*Q*| − *k* + 1)⌋} where the colors *i* are *sorted by decreasing weight w*(*i, M* (*Q*)).

We say that *R*(*Q, τ*) is a *ranked list*, as each color is ranked by its weight. Throughout this paper, we use 1-based indexing and square-bracket notation *R*[*i*..*j*] to refer to the elements from position *i* to *j* included, 1 ≤ *i* ≤ *j* ≤ |*R*|.

A common value for *τ* is, for example, 0.8, retaining documents containing at least 80% of the *k*-mers of *Q*. This query mode is used by both exact and approximate indexes in state-of-the-art methods (e.g., MetaGraph [Karasikov et al., 2020], Fulgor [Fan et al., 2024], COBS [Bingmann et al., 2019], and kmindex [Lemane et al., 2024]). Fig. 1 shows an example of a threshold-union query.

**Fig. 1.**
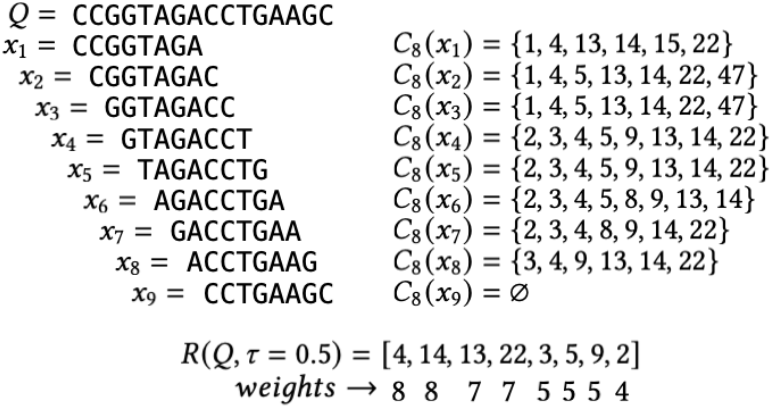
Threshold-union query example for *k* = 8 and *τ* = 0.5. The query string *Q* contains 9 *k*-mers, *x*_1_, …, *x*_9_. On the right, their hypothetical color sets. The fact that *C*_8_(*x*_9_) = ∅ means that *x*_9_ has not been indexed. The result set *R*(*Q, τ*) is computed by including all colors whose weight is at least ⌊*τ* (|*Q*| − *k* + 1)⌋ = ⌊0.5 · 9⌋ = 4, sorted by weight.

The approximate version of *R*(*Q, τ*) is indicated with

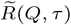 and is defined in an analogous way but over 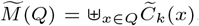. Note that, since 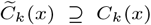 for any *x* we have that: **(1)** 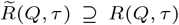 for any query *Q*, and **(2)** 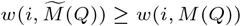 for any color *i*.

### 2.3. Rank-biased overlap (RBO)

Given two ranked lists of infinite length, Webber et al. [2010] define their *rank-biased overlap* (RBO, henceforth) as a measure of their similarity.

In this work, we exploit one particular definition: the “bounded RBO”, noted RBO@*D*, that measures the RBO of lists truncated at depths *D*. See supp. mat. for details and examples.

## 3. Related work

Since this work focuses on approximate solutions, we do not review exact indexes here. However, our experiments also report results for two exact indexes, MetaGraph [Karasikov et al., 2020] and Fulgor [Fan et al., 2024], used to determine the ground truth for the queries.

Most solutions use approximate membership query data structures, mainly indexing *k*-mers with Bloom filters [Bloom, 1970]. Practical implementations include BIGSI [Bradley et al., 2019], later enhanced by COBS [Bingmann et al., 2019]. These methods create individual Bloom filters for each document, forming final indexes as inverted matrices that interleave the filters. For a given hash value, *N* consecutive bits indicate the presence/absence of a *k*-mer across *N* documents, allowing efficient query access to all *k*-mer occurrences per sample.

Recent advancements in MetaProFi [Srikakulam et al., 2023] and kmtricks [Lemane et al., 2022] have enabled the processing of larger data volumes. Notably, kmtricks was utilized in kmindex [Lemane et al., 2024] to index hundreds of terabytes of metagenomic seawater data for the first time. This method employs the findere approach [Robidou and Peterlongo, 2021], where a unique hash function is used to reduce false positive rates by querying multiple *s*-mers per *k*-mer (with *s* ≤ *k*). This approach lowers query times by minimizing accesses to the bloom filter. Similarly, MetaProFi utilizes a chunked Bloom filter matrix with compression to significantly decrease the overall index size.

A family of methods organizes Bloom filters in a tree topology[Solomon and Kingsford, 2016; Sun et al., 2018; Harris and Medvedev, 2020; Gupta et al., 2021; Marchet and Limasset, 2023], with leaves containing Bloom filters and internal nodes storing unions of siblings. Such layout saves space by avoiding duplicating information related to *k*-mers present in a full subtree and enhances query speed by halting searches as soon as a subtree is fully determined. Nevertheless, these approaches suffer from random memory access issues that limit their query performances.

Finally, the tools Raptor [Seiler et al., 2021] and PebbleScout [Shiryev and Agarwala, 2024] are the closest solutions to our proposal. They index color sets of minimizers of *k*-mers rather than the *k*-mers themselves. For Raptor, minimizers are indexed with Hierarchical Interleaved Bloom Filters [Mehringer et al., 2022], optimizing for unbalanced input dataset sizes. In PebbleScout, color sets are indexed using minimizers of fixed length *m* = 25, and *k*-mers of length *k* = 42. While PebbleScout has been effectively used on a significant portion of the Sequence Read Archive datasets (SRA [Katz et al., 2022]), it is not open-source, preventing a direct comparison. Lastly, minimizers have also been exploited to map *k*-mers to reads [Vandamme et al., 2025].

## 4. Approximate indexing of a set of documents

The high-level idea we propose is to store color sets of minimizers, instead of color sets of *k*-mers. In other words, we take the union of color sets of *k*-mers sharing the same minimizer. For the sake of clarity, we define *C*_*m*_(*µ*) as the color set of a minimizer *µ*, i.e., *C*_*m*_(*µ*) = {*i*|*µ* ∈ *R*_*i*_}. In practice, our solution is to let 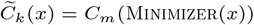.

Minimizers have some important properties that should be exploited to achieve compact space and fast query time for Problem 2.

1. *Consecutive k-mers tend to share the same minimizer*. This property allows for a drastic reduction in the size of the final index since there are fewer distinct minimizers than *k*-mers. As reviewed in Section 2.1, we expect to have approximately (*k* − *m* + 2)*/*2 times fewer (random) minimizers than *k*-mers.-----Additionally, we exploit this property when retrieving the color sets of *k*-mers: we save repeated accesses to the color set of the minimizer (i.e., we cache the set) by *streaming* through the *k*-mers of *Q*. Other solutions, like COBS, cannot exploit this streaming query pattern because every *k*-mer lookup accesses a different row of its binary matrix, resulting in a cache miss per each *k*-mer of *Q*. Also, we *skip* the query of all *k*-mers whose minimizer is not indexed.
2. *Minimizers are sufficiently well skewed, at least for reasonable large length m < k, e.g*., *for m* = 19 *and k* = 31. We thus expect to decode a short color set, whose compressed representation often takes much less than *N* bits. This is in net contrast to solutions based on uncompressed binary matrices, such as COBS, which always has to decode *N* bits.
3. *Consecutive k-mers tend to have very similar color sets* (*if not exactly the same*). As noted above, consecutive *k*-mers tend to share the same minimizer. Because of this, there is a high chance that every time we see a minimizer we also see its *k*-mers. This intuitively helps to keep under control the amount of false positives in 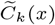 for all the *k*-mers *x* that have minimizer *µ* because *C*_*m*_(*µ*) results from the union of similar sets. On the other hand, several indexes reviewed in the previous section merge color sets of randomly-chosen *k*-mers potentially having very different sets of colors.

Some properties have been used in existing literature [Pibiri, 2022; Fan et al., 2024; Seiler et al., 2021; Vandamme et al., 2025] to address related problems; however, before this work no comprehensive solution to Problem 2 integrating all these properties was available. Notably, these properties are independent of the specific data structures used for indexing minimizers and compressing color sets, allowing for various space/time trade-offs. We detail our approach in the following sections, introducing an index named “Kaminari” that leverages these properties.

### 4.1. Index creation and description

Creation of a Kaminari index comprises four steps, represented in Figure 2.

**Fig. 2.**
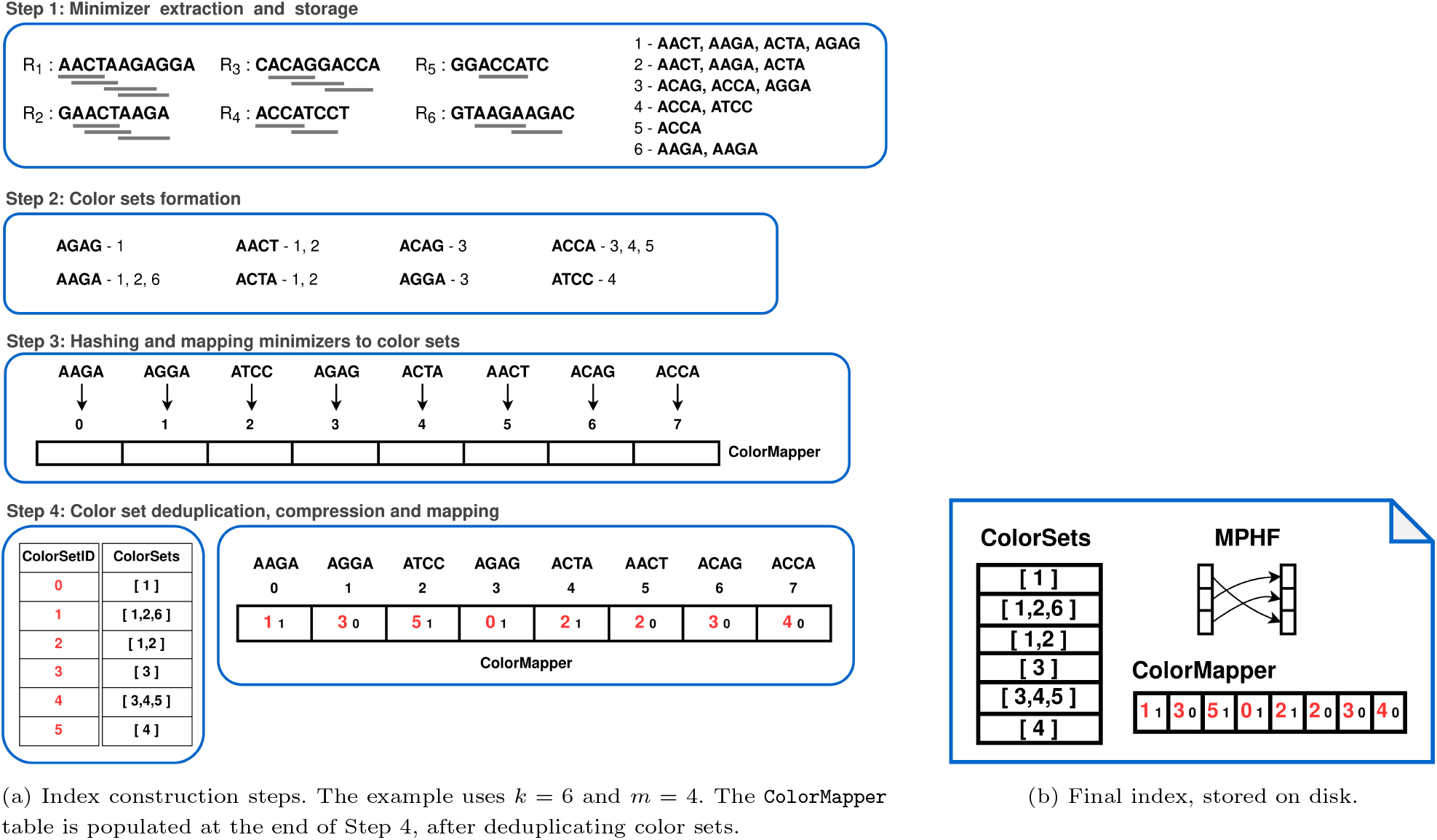
Kaminari index: (a) construction steps; (b) final representation.

#### Step 1: Minimizer extraction and storage

For each document *R*_*i*_ ∈ ℛ, all random minimizers of length *m* are computed, and their MurmurHash2 hashes^1^ are stored in an external-memory vector. This vector maintains a list of disk files (blocks) containing its elements, along with a small internal-memory buffer for the block under construction. If the total RAM used by the *N* buffers exceeds a user-defined limit, the buffers are sorted, deduplicated, and written to *N* separate blocks on disk. Let *K* ≥ *N* be the total number of blocks written.

#### Step 2: Color sets formation

We merge these *K* sorted blocks to deduplicate minimizers and construct the corresponding color sets, utilizing a classic merging strategy with log_2_ *K* parallel merges. This process also enables the incremental building of minimizer color sets, leveraging external-memory abstractions indicating each block’s logical color. Color sets can be encoded as binary vectors of *N* bits or through more advanced encodings (see below). Ultimately, we compute the distinct minimizers of ℛ and their associated color sets.

#### Step 3: Hashing and mapping minimizers to color sets

Call *S* the set of distinct minimizers of ℛ. A minimal perfect hash function (MPHF) *f* is built for *S* using PTHash [Pibiri and Trani, 2021, 2023; Hermann et al., 2024]. This function maps each minimizer to an index in a table, *ColorMapper*[1..|*S*|], created and used in last step.

#### Step 4: Color set deduplication, compression and mapping

Let *z* indicate the number of distinct color sets. Note that *z* ≤ |*S*| because different minimizers can have the same color set. We start by deduplicating color sets and assign a unique identifier 0 *< I* ≤ *z* to each distinct color set (for example, following their lexicographic order). For every *µ* ∈ *S*, we store the pair of integers (*I, F*) at position *i* = *f* (*µ*) in the *ColorMapper* table:

1. The integer *I* uniquely identifies the color set associated with *µ*.
2. The value *F* is *b*-bit integer, computed as *F* = Fingerprint(*µ*). We implement the function Fingerprint : Σ^*m*^ → [2^*b*^] as a pseudo-random hash function, that is, *F* is a pseudo-random integer in the interval [1, 2^*b*^].

Such fingerprint is used in the detection of alien minimizers (minimizers not belonging to the input collection ℛ). Suppose we query for a minimizer *α*. At query time, Fingerprint(*α*) is computed and compared against that stored in *ColorMapper*[*f* (*α*)]. If they are *not* the same, then *α* is surely alien. Otherwise, *α* is not alien with probability at least 1 − 1*/*2^*b*^. This is a folklore technique to implement a space-efficient static filter with prescribed false positive probability [Marchet et al., 2020; Broder and Mitzenmacher, 2003; Bender et al., 2018]. In the example of Fig. 2 we use *b* = 1 and these bits are displayed in small black font within the *ColorMapper* table.

It follows that the *ColorMapper* table is stored in |*S*|(*b* + ⌈log_2_ *z*⌉) bits. The color sets themselves are compressed using techniques inspired by Fulgor [Fan et al., 2024]. Specifically, each color set is classified into one of the following three categories based on its size.

- *Sparse color sets*. If the number of colors in the set is less than *N/*4, the colors are encoded using a difference-based approach: we first compute the differences between consecutive integers and then apply Elias’ *δ* encoding [Elias, 1975] to compress them.
- *Dense color sets*. When the size of the set is at least *N/*4 and at most 3*N/*4, we use a binary vector of *N* bits. A bit set at position *i* indicates that color *i* is present in the color set.
- *Very dense color sets*. Lastly, if the size of the set exceeds 3*N/*4, we encode only the absent document identifiers using the same difference-based approach as for the sparse sets.

All the compressed representations of the color sets are concatenated in a single bitvector called *ColorSets* in Fig. 2. The starting positions of the color sets are kept in a separate list, so that we can accessed the compressed representation of the *i*-th color set for any 0 *< i* ≤ *z*. Since this list is monotone by construction, we compress it using the Elias-Fano encoding [Elias, 1974; Fano, 1971].

To sum up, the Kaminari index consists of the following three components: (1) the MPHF *f*, (2) the *ColorMapper* table, (3) and the compressed color sets.

### 4.2. Queries

The crux of computing 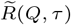 is how 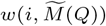 is determined efficiently *without* explicitely materializing the set 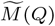, i.e., without taking the multi-set union of all the colors sets for all the *k*-mers of *Q*. As it is clear, we need to only consider the color sets for the minimizers of the *k*-mers of *Q, Z* = Minimizers(*Q*). Since each *µ* ∈ *Z* appears in *Q* for *w*(*µ, Z*) times by definition, the weight of the color *i* in 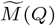 is just the sum of the weights *w*(*µ, Z*) for all color sets *C*_*m*_(*µ*) where *i* appears. In formal terms, 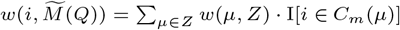, where I[*E*] is the indicator variable for the event *E*, that is, I[*E*] = 1 if *E* is true and 0 otherwise. This allows us to efficiently compute 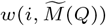 in an incremental way (Algorithm 1). We initially set 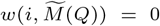 and, when scanning *C*_*m*_(*µ*), *w*(*µ, Z*) is summed to 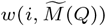 if *i* ∈ *c*_*m*_(*μ*).

It remains to explain how the set *C*_*m*_(*µ*) is retrieved from the index (Algorithm 2). First, *p* = *f* (*µ*) is computed and the pair (*j, F*) = *ColorMapper*[*p*] is retrieved. If *F* matches the fingerprint of *µ*, then we set *C*_*m*_(*µ*) = *ColorSets*[*j*]; otherwise *C*_*m*_(*µ*) = ∅.

### 4.3. False positives

With the proposed scheme, there are two different sources of false positives when computing 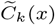 as *C*_*m*_(Minimizer(*x*)).

1. The first cause is that the color set of a *k*-mer is a subset of the color set of its minimizer, i.e., *C*_*k*_(*x*) ⊆ *C*_*m*_(Minimizer(*x*)). Hence using *C*_*m*_(Minimizer(*x*)) as an approximation for *C*_*k*_(*x*) clearly introduces false positives, i.e., spurious colors that are due to *C*_*k*_(*y*) ⊂ *C*_*m*_(Minimizer(*y*)) for any other *k*-mer *y* ≠ *x* such that Minimizer(*y*) = Minimizer(*x*). This effect is exacerbated when the *k*-mer is absent but the minimizer is present. In this case, all the colors in the returned color set are false positives.
2. The second cause is that the MPHF *f* — by definition — cannot detect whether a minimizer has been indexed or not. For an alien minimizer *µ*, we remark that *f* (*µ*) can be any integer in {1, …, |*S*|}. This implies that Kaminari always returns a color set, even when an alien minimizer is queried. In such cases, the correct color set would be the empty set and therefore all elements of the returned set must be considered false positives.

To mitigate this effect, we use the *b*-bit fingerprint *F* stored along with the identifier of the color set of each minimizer. As already noticed, Fig. 2 shows an example with *b* = 1, so that we reject alien minimizers for approximately 50% of the time.

#### Algorithm 1

The threshold-union query for a sequence *Q* and parameter *τ*, as supported by Kaminari. The algorithm computes the ranked list 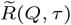 as described in Section 2.2, i.e., by returning all colors *i* such that 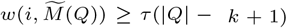. For ease of notation, we let 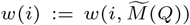 and 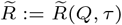 in the pseudocode.

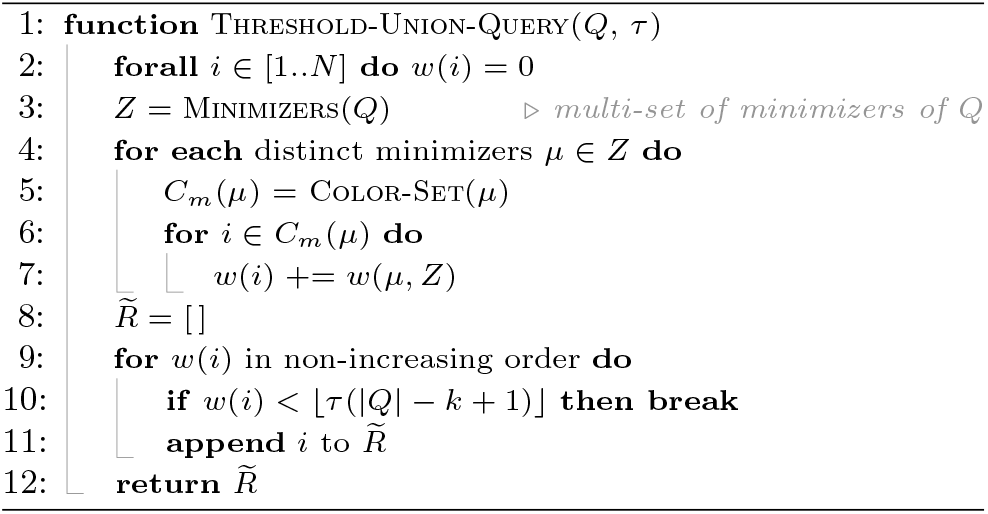

#### Algorithm 2 The retrieval of the color set of the minimizer *µ* from a Kaminari index.

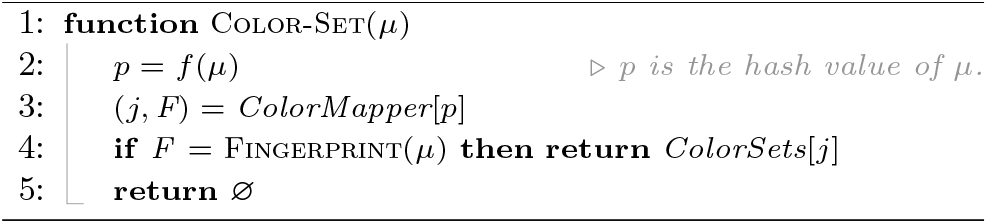

Fig. 3 shows an example of these two effects, using the same query string *Q* from Fig. 1. Note how, for example, color 19 is a false positive for the first of the two reasons described above: it belongs to both *C*_4_(*µ*_1_) and *C*_4_(*µ*_3_) and has a weight of 6 *>* ⌊*τ* (|*Q*| − *k* + 1)⌋ = 4. This means that minimizers CCGG and ACCT appear in *R*_19_, leading to color 19 to be associated to all *k*-mers having these two minimizers (in the example, *x*_1_ and *x*_4−8_). On the other hand, as an example of alien minimizer lookups, we reconsider the example from Fig. 1. There, *k*-mer*x*_9_ does not appear in any document of ℛ. Let us further assume that this so because its minimizer, AAGC, is an alien minimizer. While the returned color set *C*_4_(*µ*_4_) can be any of the indexed color sets, the fingerprint matching strategy correctly identifies it as an alien minimizer. Note how this prevents the color 8 to gain a weight of 8.

**Fig. 3.**
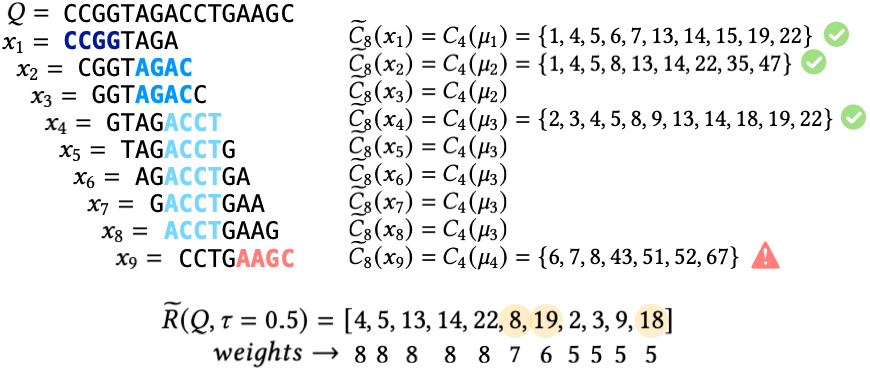
The same example discussed in Fig. 1 but in the approximate setting presented in Section 4, i.e., assuming that *C*_*k*_(*x*_*i*_) = *C*_*m*_(*µ*_*j*_) for a *k*-mer *x*_*i*_ whose minimizer is *µ*_*j*_. The lexicographic minimizer of length *m* = 4 of each *k*-mer is indicated in bold font; we have *µ*_1_ = CCGG, *µ*_2_ = AGAC, *µ*_3_ = ACCT, and *µ*_4_ = AAGC. We assume that *µ*_1−3_ match the fingerprint stored in the index (green check mark), whereas *µ*_4_ does not (red warning) and it is correctly labeled as alien. Compared to the exact result set *R*(*Q, τ*) = [4, 14, 13, 22, 3, 5, 9, 2] from Fig. 1, the set *R*(*Q, τ*) contains three false positives, {8, 19, 18} (highlighted in yellow), and the ranking of the reported colors is different.

The expectation is that false positives do not occur in more than *τ* · 100% of the color sets of the *k*-mers of *Q*, and are thus not reported in the final result 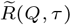.

## 5. Results

Kaminari is written in `C++` and was compiled with `gcc` 14.2.0 for the experiments illustrated here. Reproducibility scripts, including scripts to download all tested collections and to build and query the indexes can be found at github.com/vicLeva/benchmarks kaminari.

We report the performance of Kaminari in terms of disk size, query time, and resource usage during construction. Effectiveness is measured using the RBO similarity, as stated in Section 2. For all experiments, we considered canonical *k*-mers, with *k* = 31 and *τ* = 0.8 for queries, motivated by results presented Figure 4 in appendix.

**Fig. 4.**
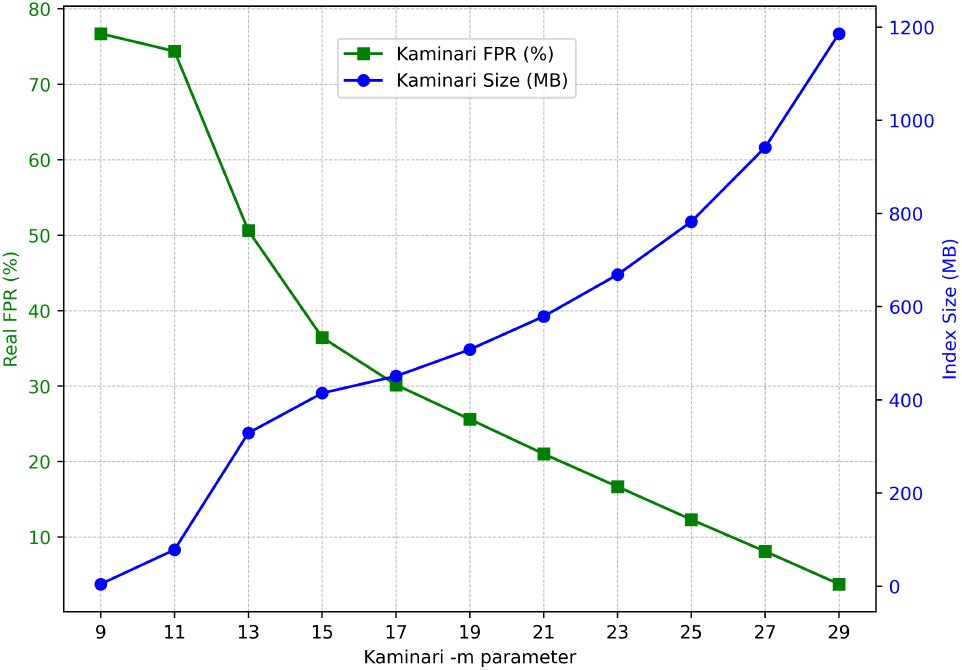
Kaminari index size and FPR measured on positive queries by varying the m parameter when built on the Ecoli dataset. k is fixed to 31.

### Hardware

All experiments were conducted on a GenOuest platform on a node with 4 × 8 cores Xeon E5-2660, clocked at 2.20 GHz and with 1.5 TB of memory, running CentOS Linux 7 (Core) server with a 64-bit architecture (kernel version 3.10.0).

### Datasets

We consider five collections characterized by different file counts, average lengths, and internal similarity. Basic statistics are provided in Table 1.

**Table 1.**
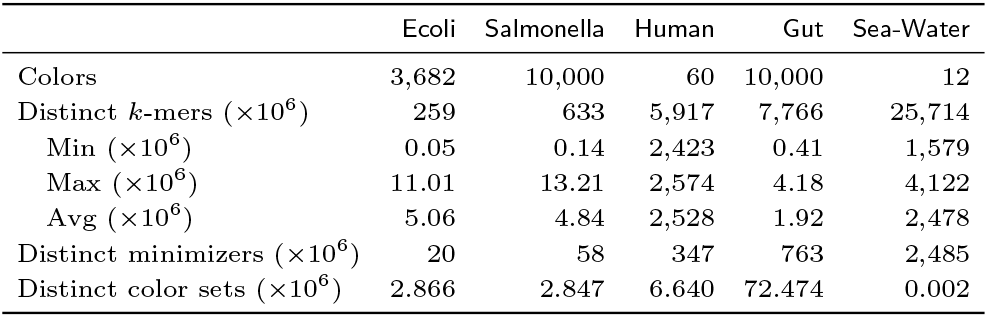
Some basic, approximate, statistics for the tested collections for *k* = 31 and *m* = 19 as minimizer length.

- Ecoli: 3,682 *E. Coli* genomes.
- Salmonella: 10,000 *S. Enterica* genomes.
- Human: 60 whole human genomes (30 paternal haplotypes; 30 maternal).
- Gut: 10,000 Gut metagenome-assembled genomes (MAG).
- Sea-Water: A collection of 12 metagenomics non-assembled samples from sea water.

### Competitors

We compared the performance of Kaminari against 2 exact tools: Raptor and MetaGraph alongside with 3 approximate tools: COBS, kmindex and Raptor. More details about these tools in Appendix. Rambo is excluded due to its index size being up to twice that of COBS, and HowDe-SBT is not included as its construction and queries are significantly slower than those of COBS [Bingmann et al., 2019]. Among the tested tools, Fulgor and MetaGraph are exact solutions, yielding no false positive matches. They serve as baselines for evaluating the tradeoffs of approximate solutions. Additionally, we compared the exact rankings from Fulgor with those from non-exact tools using the RBO measure to assess the latter’s effectiveness.

All indexes were evaluated using the `C++` implementations provided by the authors. Default parameters were employed unless otherwise noted, and tools were provided 32 threads to operate. We report in Appendix the tested tool versions and the exact parameters used.

### 5.1. Efficiency

#### Index size

Table 2 shows the disk size of each index and raw data. Kaminari consistently produces the smallest index, often orders of magnitude smaller, with the exception of MetaGraph, which utilizes the BOSS [Bowe et al., 2012] data structure. Results show that Kaminari performs well across various data types, including assembled bacterial and eukaryotic genomes, MAG genomes, and complex raw data. Although MetaGraph generates compact indexes, it suffers from significantly longer query times, as shown in the next section.

**Table 2.** Comparisons of index size on disk and raw data size (compressed with gzip). All values are in gigabytes. Color code is green: *<* 3 × *best*, orange: *<* 10 × *best*, red otherwise.

|  | Ecoli | Human | Salmonella | Gut | Sea-Water |
| --- | --- | --- | --- | --- | --- |
| Raw data size (gzip) | 5.52 | 48.53 | 14.17 | 5.72 | 38.79 |
| Kaminari | 0.50 | 1.13 | 0.81 | 4.78 | 4.51 |
| Fulgor | 1.35 | 4.67 | 2.28 | 12.23 | 37.54 |
| COBS | 7.01 | 64.07 | 17.59 | 6.95 | 109.50 |
| kmindex | 2.91 | 11.02 | 9.49 | 5.43 | 4.51 |
| Raptor | 2.71 | 17.75 | 7.37 | 4.57 | 8.72 |
| MetaGraph | 0.33 | 3.09 | 0.60 | 3.88 | 11.05 |

#### Query performance

The second key result is Kaminari’s query speeds, reported Table 3 (and Table 8 in Appendix).

**Table 3.** Time and peak memory usage for 50,000 positive queries (1000 base pairs each), using *τ* = 0.8. See Table 2 for color code.

|  | Ecoli |  | Human |  | Salmonella |  | Gut |  | Sea-Water |  |
| --- | --- | --- | --- | --- | --- | --- | --- | --- | --- | --- |
|  | seconds | GB | seconds | GB | seconds | GB | seconds | GB | seconds | GB |
| Kaminari | 7 | 0.6 | 3 | 1.2 | 22 | 1.2 | 10 | 4.8 | 10 | 4.6 |
| Fulgor | 21 | 1.5 | 8 | 4.7 | 62 | 2.4 | 16 | 12.3 | 29 | 37.6 |
| COBS | 80 | 7.4 | 103 | 64.1 | 166 | 18.6 | 52 | 7.9 | 121 | 109.5 |
| kmindex | 88 | 24.2 | 19 | 2.5 | 328 | 60.3 | 91 | 61.2 | 11 | 3.1 |
| Raptor | 5 | 2.8 | 9 | 17.8 | 12 | 7.4 | 5 | 4.6 | 5 | 8.8 |
| MetaGraph | 3619 | 0.5 | 262 | 3.1 | 13065 | 0.8 | 200 | 4.0 | 27 | 11.1 |

These results show the total elapsed time to perform 50,000 queries for each dataset, each query consisting of a sequence composed of 1,000 bases. Results with different query lengths are proposed in the companion repository. In case of so-called “*positive queries*” (Table 3), these queried sequences were randomly taken from the documents used to build the index. On the other hand, “*negative queries*” (Table 8 in Appendix) are random sequences absent from the documents.

With the exception of kmindex, RAM usage reflects the size of the index for all tools. kmindex does not load the full index in memory. Additionally, it is optimised so that parts of the index are mapped to RAM and remain in RAM as long as memory is available, limiting the number of disk accesses. This explains its higher RAM usage with 50,000 queries.

The MetaGraph results show prohibitive query times, up to≈ 600 times slower than Kaminari when performing positive queries. Raptor also shows good time performances, however, its index sizes and, consequently its RAM usage is up to an order of magnitude bigger than Kaminari ones.

Overall, Kaminari results are always among the best ones, if not the best. In contrast, all other tested solutions have at least one instance where query time or RAM usage is an order of magnitude greater than that of Kaminari.

#### Construction performance

Table 9 (Appendix) shows that the construction results for Kaminari are good, if not the best, for genomic datasets (Ecoli, Human, Salmonella). For more complex datasets like Gut and Sea-Water, tools such as kmindex perform better due to their specialized construction algorithms. We propose two construction strategies: the default method described in Chapter 4.1 and a second one enabled by the `-metagenome`option. This option, intended for datasets with over 128 documents composed solely of metagenomic data, reduces redundancy *intra* and *inter* documents, resulting in a high number of minimizers and shorter color sets. Therefore, in this case, we encode color sets as lists of integers rather than using binary encoding.

### 5.2 Effectiveness with default parameters

In this section we provide effectiveness results while using all tools with their default parameters. We believe that these information are useful from a user perspective. However, these comparisons might appear unfair as, under these conditions, FPRs and index sizes are not directly comparable between different tools. In Section 5.3 and Section 5.4, we propose results fixing one of these two parameters at a time.

#### False positives evaluation

As we use *τ* = 0.8, a document is defined as a false positive if 80% or more of the *k*-mers of the query sequence are reported as present whereas an exact index would correctly mark it as absent.

Table 4 shows that Kaminari and COBS produce similar results, with distinct false positive sources: minimizers and bloom filters. In contrast, Raptor accumulates imprecision from both, leading to more false positives. The *Findere* method allows kmindex to attain the best precision, despite significantly slower query times.

**Table 4.** False positive rates (%), using default parameters for all tools (recalling that index sizes for COBS, kmindex and Raptor are≈ 3 to 21 times bigger than for Kaminari).

|  | Ecoli | Human | Salmonella | Gut |
| --- | --- | --- | --- | --- |
| Kaminari | 25.61 | 1.24 | 32.08 | 5.83 |
| COBS | 25.83 | 0.87 | 31.64 | 5.84 |
| kmindex | 5.81 | 1.06 | 5.00 | 0.60 |
| Raptor | 31.14 | 1.27 | 36.67 | 7.97 |

For negative (random) queries, we observed that all tested tools reported 0 false positives.

#### RBO results

False positives arise from over-estimations that cause a document to incorrectly exceed the threshold *τ*. We think that the real impact for the user is better highlighted by the RBO metric.

We compared Kaminari answers to the ground truths provided by Fulgor by using the RBO metric to estimate the impact of the false positives in real-life applications in which top hits are the most informative. For every query, we computed RBO(*R*(*Q, τ*), 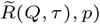, with *R*(*Q, τ*) being the ordered list of documents generated by Fulgor and 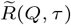 the one generated by non-exact tools, including Kaminari. The RBO value depends on a parameter *p*. We explain in appendix how this parameter was determined.

While RBO can be applied to very short lists (e.g., size *<* 10), it is not very informative in such case due to limited overlap measurement opportunities. Therefore, we present Table 5 RBO metrics only for lists of size ≥ 10. We did not include Sea-Water results as the dataset contains only 12 samples. Similarly, Raptor is also excluded due to its lack of ranked output, rendering RBO evaluation infeasible.

**Table 5.**
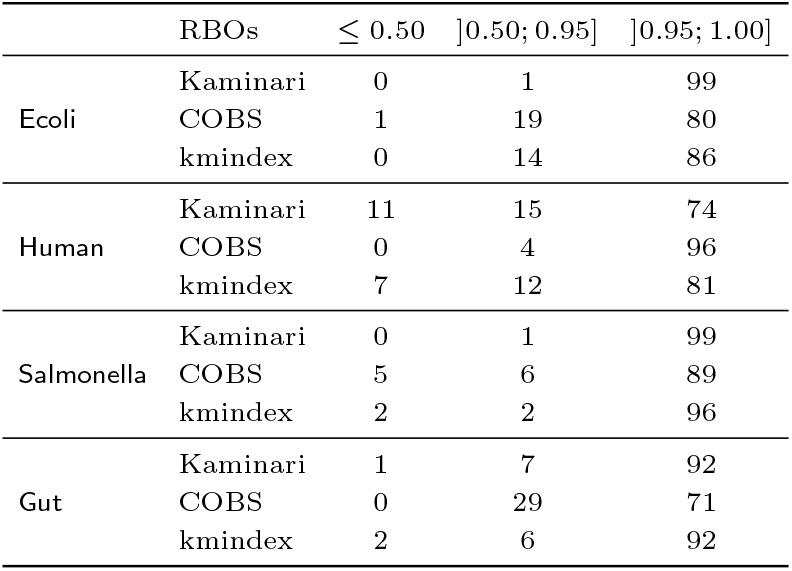
RBO values distribution for positive queries, for truth lists of size ≥ 10. In each column, we report the proportion (%) of queries whose RBO value are in the specified range. Full results are provided Figure 5 in appendix.

All datasets and tools show a significant skew toward high RBO values, with most queries near an RBO of 1. This suggests that tools addressing Problem 2 produce rankings closely resembling the ground truth across various datasets. Except for the Human dataset, Kaminari consistently achieves the highest RBO values across biological domains, demonstrating its robustness and effectiveness in capturing meaningful rankings. In the case of the Human dataset, the RBO metric encounters limitations due to the extreme similarity of samples, leading to subtle variations in query answers. Consequently, minor discrepancies from non-exact methods can alter ranking results. Nevertheless, as shown previously, the false positive rate for this dataset remains low, at 1.24% for Kaminari, thus preserving the biological significance of the findings.

### 5.3. Effectiveness fixing the index sizes

In this section, for all tested tools, we fixed the index size to the one obtained by Kaminari. In this configuration, false positive rates, shown Table 6, highlight that on genomic data, for the same disk size budget, Kaminari generates the least amount of false positives. On the metagenomic Gut dataset, kmindex performs better despite being ≈ 9 times slower to query. RBO results (Figure 6 in Appendix) additionally show that in this setup, Kaminari provides more precise ranking of the results, for any of the considered dataset.

**Table 6.** False positive rates (%), using the same index size for all tested tools.

|  | Ecoli | Human | Salmonella | Gut |
| --- | --- | --- | --- | --- |
| Kaminari | 25.61 | 1.24 | 32.08 | 5.83 |
| COBS | 76.67 | 1.92 | 59.27 | 10.57 |
| kmindex | 76.59 | 1.92 | 59.21 | 0.79 |
| Raptor | 62.91 | 1.92 | 58.11 | 7.99 |

### 5.4. Effectiveness fixing the FPR

So far, every tool has been used and measured with default parameters. In Table 7, we adjusted the parameters of the approximate tools in order to achieve 10% of false positives in queries. Human and Sea-Water datasets were not included because 10% could not be reached due to the data redundancy and documents number. Results are similar to Table 2 in the sense that Kaminari provides the best results, except for metagenomes for which it nevertheless remains competitive.

**Table 7.** Index Size (GB), while empirically producing results with 10% of false positives. With *k* = 31, Raptor could not reach 10% of false positives for Salmonella. More details about the parameters used can be found in Appendix.

|  | Ecoli | Salmonella | Gut |
| --- | --- | --- | --- |
| Kaminari | 0.94 | 1.73 | 3.44 |
| COBS | 20.29 | 65.28 | 4.52 |
| kmindex | 2.20 | 6.51 | 2.71 |
| Raptor | 15.76 | NA | 2.93 |

**Table 8.** Total elapsed time (seconds) and peak memory usage (GB) for 50,000 negative queries (1000 base pairs), using *τ* = 0.8. See Table 2 for color code.

|  | Ecoli |  | Human |  | Salmonella |  | Gut |  | Sea-Water |  |
| --- | --- | --- | --- | --- | --- | --- | --- | --- | --- | --- |
|  | seconds | GB | seconds | GB | seconds | GB | seconds | GB | seconds | GB |
| Kaminari | 2 | 0.6 | 3 | 1.2 | 2 | 0.9 | 10 | 4.8 | 10 | 4.6 |
| Fulgor | 1 | 1.5 | 8 | 4.7 | 2 | 2.3 | 13 | 12.3 | 26 | 37.6 |
| COBS | 56 | 7.4 | 98 | 64.1 | 55 | 18.6 | 52 | 7.9 | 118 | 109.5 |
| kmindex | 41 | 24.2 | 17 | 2.5 | 110 | 60.3 | 84 | 61.2 | 11 | 3.0 |
| Raptor | 1 | 2.8 | 10 | 17.8 | 4 | 7.4 | 3 | 4.6 | 5 | 8.8 |
| MetaGraph | 7 | 0.4 | 18 | 3.2 | 11 | 0.7 | 23 | 4.0 | 35 | 11.1 |

**Table 9.** Index construction time (h:mm:ss) and peak memory usage (GB). See Table 2 for color code.

|  | Ecoli |  | Human |  | Salmonella |  | Gut |  | Sea-Water |  |
| --- | --- | --- | --- | --- | --- | --- | --- | --- | --- | --- |
|  | h:mm:ss | GB | h:mm:ss | GB | h:mm:ss | GB | h:mm:ss | GB | h:mm:ss | GB |
| Kaminari | 0:02:01 | 10.12 | 0:08:53 | 46.00 | 0:11:31 | 72.47 | 0:50:09* | 96.96* | 1:38:59* | 294.77* |
| Fulgor | 0:08:36 | 16.45 | 0:38:44 | 219.10 | 0:14:45 | 21.84 | 0:53:04 | 50.17 | 2:17:00 | 153.70 |
| COBS | 0:02:19 | 6.09 | 1:02:43 | 64.07 | 0:26:27 | 35.21 | 0:04:05 | 6.38 | 1:13:36 | 27.38 |
| kmindex | 0:07:07 | 3.71 | 0:37:01 | 8.12 | 0:38:36 | 3.38 | 0:19:29 | 10.24 | 0:08:48 | 43.74 |
| Raptor | 0:02:03 | 4.62 | 1:20:36 | 833.71 | 0:05:57 | 9.23 | 0:02:59 | 8.91 | 0:44:19 | 354.71 |
| MetaGraph | 0:44:50 | 141.65 | 3:34:21 | 284.46 | 2:39:49 | 257.08 | 2:17:54 | 148.04 | 2:36:16 | 256.18 |

\* using the Kaminari *-metagenome* option

## 6. Conclusions and future work

In this work, we introduced Kaminari, a novel approximate approach for indexing sets of genomic sequences. By leveraging the properties of *k*-mer minimizers, Kaminari achieves significant improvements over traditional Bloom filter-based solutions in terms of both memory efficiency and query performance. We believe this approach will enable the creation of indexes for massive datasets, in the terabyte regime, while significantly reducing query time.

Some tools are more suited to indexing and querying set of closely related genomes (e.g., Fulgor [Fan et al., 2024]) while others are better tailored for complex non-assembled datasets (e.g., kmindex [Lemane et al., 2024]). On genomic datasets Kaminari always generates the smallest index (with the notable example of MetaGraph [Karasikov et al., 2020, 2022] which, however, suffers from prohibitive query times). On metagenomic dataset, kmindex achieves the best FPR for fixed index size, despite being approximately 9× slower to query. Overall, Kaminari consistently ranks as one of the fastest tools across all data types, generating the smallest indexes (or the lower FPR), often achieving the top performance and providing qualitative rankings of results. This robustness represents a key advantage when indexing heterogeneous and/or poorly characterized datasets.

This work pioneers the use of Rank-Biased Overlap (RBO) metric to evaluate similarity between ranked lists of results. Unlike traditional false positive measurements, RBO quantifies how approximation impacts result ordering — a crucial metric for assessing bias in non-exact indexing methods. We propose this evaluation framework as a new standard for measuring approximate indexing quality. While Kaminari may generate some false positives answers, RBO results showed that the impact on the ranking of colors is small. For instance, results on Ecoli show that 99% of queries present a RBO higher than 0.9, indicating the reliability of the top results.

Future work will study the use of partitioned indexes in external-memory for scaling up to even larger collections, and *repetition-aware compression* [Campanelli et al., 2024] to compress the color sets even further.

## 9. Appendix

### Rank-biased overlap (RBO)

Given two ranked lists of infinite length, *A* and *B*, Webber et al. [2010] define their *rank-biased overlap* (RBO, henceforth) as a measure of their similarity. Let *X*_*d*_ = |*A*[1..*d*] ∩ *B*[1..*d*]| be the overlap between the prefixes of length *d*. For a given parameter 0 *< p <* 1, the similarity is defined as

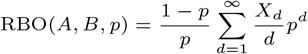

where *X*_*d*_*/d* is a measure of “agreement” between the prefixes of length *d*. Clearly, the similarity lies in [0, 1]: a value of 0 means that the two rankings are disjoint and 1 means that they are identical.

#### Bounding RBO

Although RBO is defined over infinite-length ranked lists, the summation must be truncated at a given depth *D* in practice. Call RBO@*D* (read “RBO at depth *D*”) the truncated RBO value. It is easy to see that RBO@*D* provides a lower bound to the true value of RBO, i.e., RBO *>* RBO@*D*, if RBO@*D >* 0. However, Webber et al. [2010] derive a tighter lower bound as

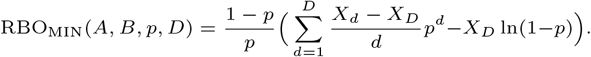

In this paper, we use the above formula with the largest possible *D*, that is *D* = min{|*A*|, |*B*|}. We set *A* = *R*(*Q, τ*) and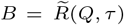. Since 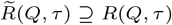,we have *D* = |*A*|.

#### Determining *p* for RBO computations

The choice of *p* is of utmost importance for RBO as it influences the result. [Webber et al., 2010] derived a formula to retrieve the weight of a prefix of the lists according to the bias parameter *p*:

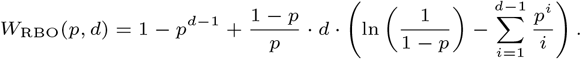

As an example, *W*_RBO_(0.85, 17) = 0.9846 means that the first 17 elements of the lists will weight for 98.46% of the RBO value. For positive queries, the length of the lists can vary from 1 to *N*. With such a variability, we made the choice to adapt *p* according to the length of the lists. More precisely, for every query, we fixed *p* so that *W*_RBO_(*p*, ⌈0.1 · |*R*(*Q, τ*)|⌉) ≈ 0.9. In other words, we want the top 10% of the list’s elements to explain 90% of the RBO value. To determine *p* for a given *d* so that *W*_RBO_(*p, d*) ≈ 0.9, we can exploit the following fact.

##### Fact 1

For fixed *d, W*_RBO_(*p, d*) is decreasing as *p* increases.

*Proof*. We show that 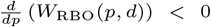. For the first term 1 − *p*^*d*−1^, we have 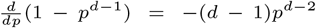. Call 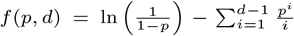 and consider the second

Term 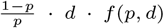. We have 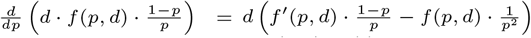. Now, we compute 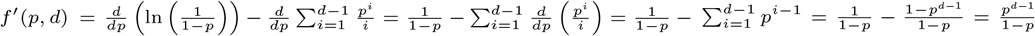. Hence by simplifying, we obtain that 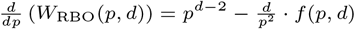.To conclude, we show that 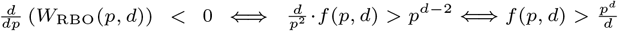. Recall that, for 0 *< p <* 1, the Taylor expansion of the logarithm is 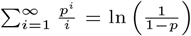

We can rewrite 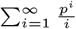 as 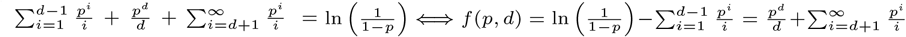 .The latter quantity is clearly larger than 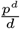 as 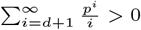.

Thus, we simply calculate the function *W*_RBO_(*p, d*) for increasing *p* ∈ (0, 1) and return the first value of *p* such that *W*_RBO_(*p, d*) ≈ 0.9. In practice, we consider the values *W*_RBO_(*i* · *ε, d*) for *i* = 1, …, ⌈(1 − *ε*)*/ε*⌉ + 1 and return the first (i.e., largest) value *p* = *i* · *ε* for which *W*_RBO_(*p, d*) *<* 0.9. The smaller *ε*, the better the approximation.

#### Interpretation and effectiveness

The parameter *p* affects the “importance” given by the top-ranked elements and has a natural probabilistic interpretation. In fact, *p* can be regarded as the probability that a user considers the next element in the ranking: a low *p* value indicates that the user is satisfied with the top results only; vice versa, a high value indicates the user’s willigness to consider more elements down in the ranking.

The RBO measure is particularly useful in scenarios where the *order* of the elements in the lists matters *more than their presence* lower down the ranking. For example, 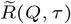 could contain a large amount of false positives but appearing at low rank positions, while the top-ranked colors could indeed be identical to those in *R*(*Q, τ*). A high RBO value thus indicates that — even in the presence of false positive matches — the ranking produced by the proposed index aligns closely with the true ranking. This ensures that the most relevant documents still appear at the top, preserving the overall utility of the retrieval process despite approximation errors.

**Example**. Consider the ranked list *R* = [4, 14, 13, 22, 3, 5, 9, 2] from Fig. 1 and 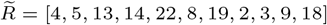 from Fig. 3 (computed using a method that we will explain in Section 4). We have 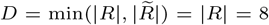. Using *p* = 0.5, we have an RBO similarity of 0.801739 and RBO_MIN_ is 0.804121. With a lower *p*, for example *p* = 0.3, the two scores are higher and more similar to each other: 0.867608 and 0.867650, respectively.

Tool versions and used parameters

- Kaminari: commit `a323d57`, parameters: `-m 19`
- Fulgor: commit `5ac5699`, parameters: `-m 19`.
- COBS: commit `2fbb044`, parameters: `--compact-construct`,
- kmindex: version `0.5.2`, parameters: `-k 25`, `-z 6`. To query 31-mers, kmindex considers 25-mers using the findere approach [Robidou and Peterlongo, 2021].
- Raptor: version `3.0.1`, parameters: `--kmer 19 --window 31`
- MetaGraph: version `0.3.6` (commit `5c2a12b`). In particular, we built the indexes following the methodology from [Fan et al., 2024] (reproducible with the workflow available at https://github.com/theJasonFan/metagraph-workflows): the indexes use the “relaxed row-diff” BRWT data structure [Karasikov et al., 2020], which is the most compact variant of MetaGraph.

About Kaminari, the choice of *m* impacts the size and the false positive rate of the index. Figure 4 shows the trade-off between performance and precision. We think *m* = 19 is a reasonable choice considering our needs.

#### 9.3. Additional results

##### 9.3.1. Performances for negative queries

Similar conclusions apply to those drawn in the main text to negative queries (Table 8), with two notable differences. Firstly, Fulgor excels in quickly detecting the absence of queried *k*-mers due to the SSHash data structure [Pibiri, 2022]. Secondly, MetaGraph queries do not experience the significant computation time issues seen with positive queries.

##### 9.3.2. Time and RAM for building indexes

Table 9 provides index construction times and peak memory usage accross tools and datasets.

##### 9.3.3. RBO distribution, full results

Figure 5 shows RBO results distribution, when using default parameters of tested tools. This is an extension of results presented Table 5.

**Fig. 5.**
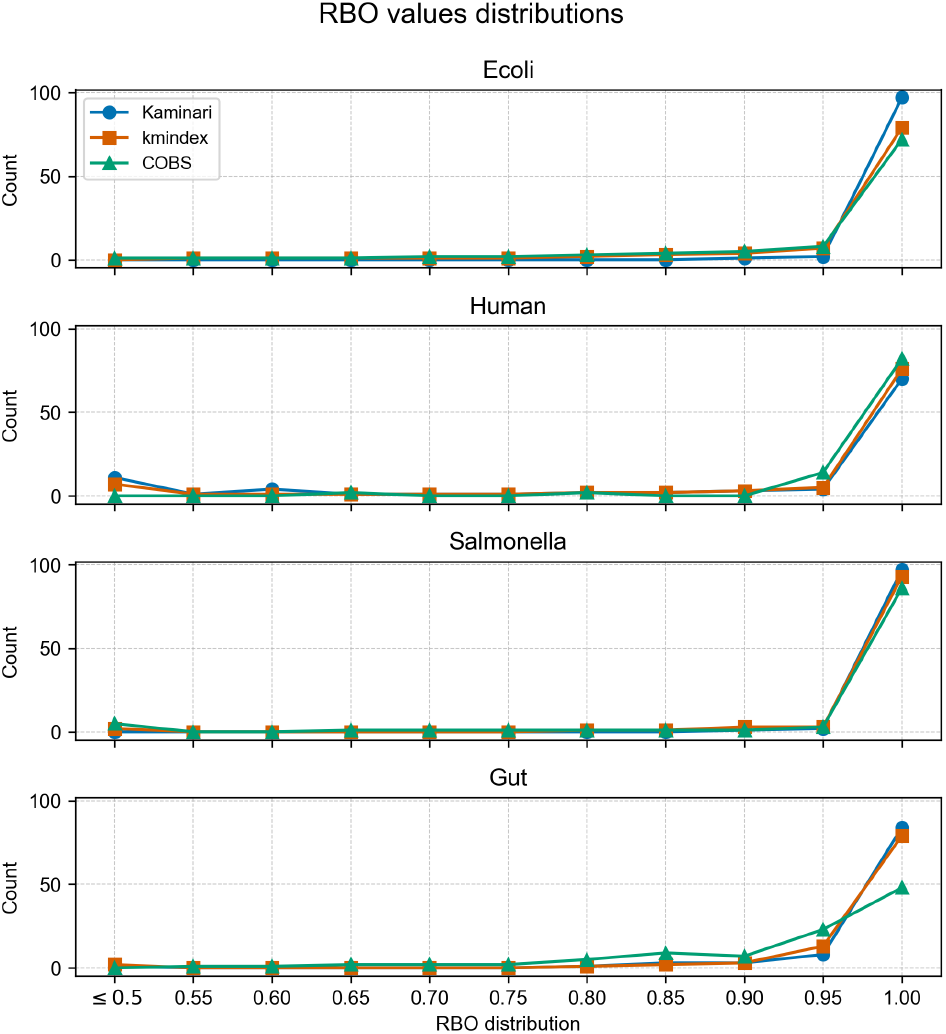
RBO values distribution for positive queries, for truth lists of size ≥ 10. Each point shows the sum of the percentage of queries from its *x* value (included) to the previous one (excluded). The leftmost point sums the percentage of queries whose RBO values are in [0, 0.5].

##### 9.3.4. RBO distribution, using equal index sizes for all tools

Figure 6 shows RBO distribution while using the same index size for all tested tools.

**Fig. 6.**
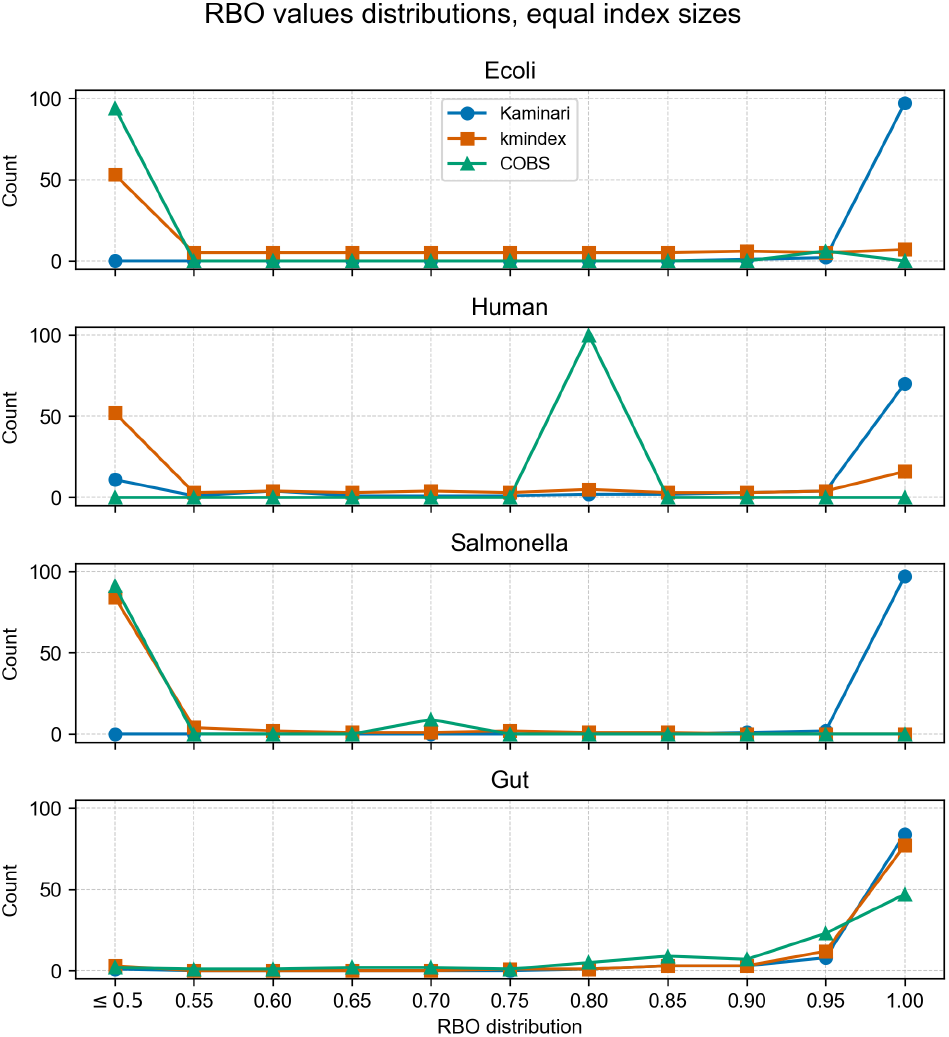
RBO values distribution for positive queries, for truth lists of size ≥ 10. In this setup index sizes are identical and are equal to Kaminari’s one. See Figure 5 for details this result representation.

##### 9.3.5. Raptor’s trade-off

Despite proposing ranked results, Raptor appears to be a serious competitor when it comes to performances (index size, query speed). In fact, it can reach Kaminari’s index size under certain parameters. Although as shown in Figure 7, when Raptor has the same index size for Ecoli than Kaminari (blue curves), its false positive rate is almost twice as big (green curves). In fact, with any parameters, Kaminari acts like a lower bound for both size and FPR for Raptor.

**Fig. 7.**
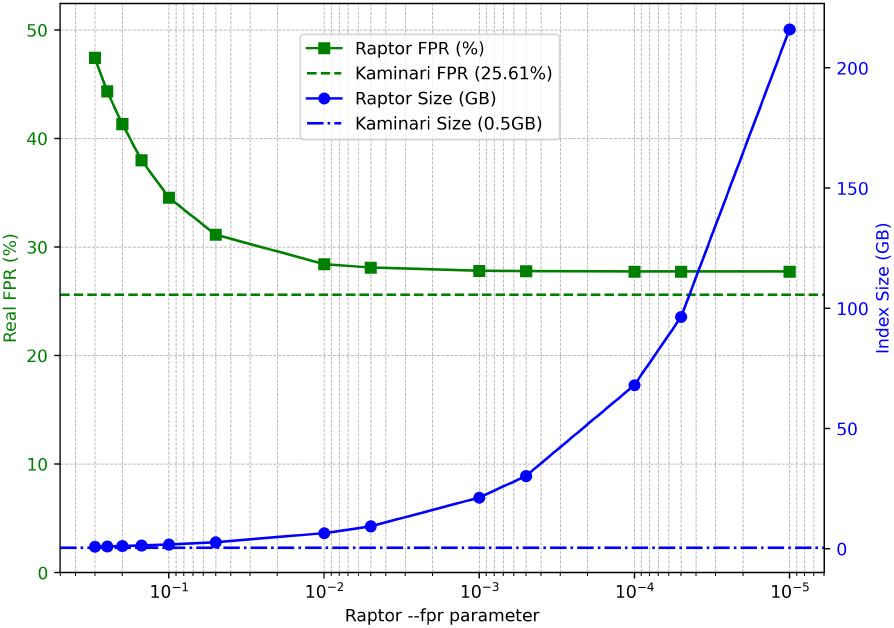
Raptor index size and FPR measured on positive queries by varying the FPR parameter when built on the Ecoli dataset. Kaminari index size and FPR are parameter independent and are indicated by dashed lines.

##### 9.3.6. Parameters

Table 6 and Table 7 present results where we tweaked some parameters to reach certain results. In any case, we kept *k* = 31. For COBS and kmindex, we changed the parameters `--false-positive-rate` and `--bloom-size`, respectively, for both experiments. About Raptor, in Table 6, we kept *m* = 19 (called `--kmer-size` in Raptor) as it corresponds to Kaminari’s *m* value, then we modified the `--false-positive-rate` parameter. In Table 7, as we modified *m* in Kaminari, we also did in Raptor. Thus, in this second experiment, we tweaked `--kmer-size` and `--false-positive-rate` for Raptor. Note that for Salmonella, even with `--kmer-size 30` and `--false-positive-rate 0.0001`, a FPR of 10% could not be reached. Exact parameters are summarized in the companion repository.

## Footnotes

1 Assuming |Σ| = 4 and *m* ≤ 32 as in our experiments, we use 64-bit hashes.

